# Editing the *CsDMR6* Gene in Citrus Results in Resistance to the Bacterial Disease Citrus Canker

**DOI:** 10.1101/2022.02.14.480359

**Authors:** Saroj Parajuli, Heqiang Huo, Fred G. Gmitter, Yongping Duan, Feng Luo, Zhanao Deng

## Abstract

Citrus is one of the most important fruit crops in the world. Citrus production worldwide faces challenges from devastating bacterial diseases, including citrus canker caused by *Xanthomonas citri* ssp. *citri* (Xcc). Improving citrus resistance to citrus canker and other major bacterial diseases has been a top priority in citrus biotechnology. Disabling disease susceptibility genes has emerged as a novel, promising approach to engineering disease resistance. The bottleneck for applying such an approach has been the identification of proper disease susceptibility-related genes in citrus. Here we show the first successful case of editing the *CsDMR6* gene in citrus and obtaining strong resistance to citrus canker in six mutants in two citrus cultivars, ‘Duncan’ grapefruit and Carrizo citrange. Multiple types of deletions and insertions were induced in *CsDMR6*, resulting in frameshift of its coding region and presumably loss of gene function. The mutation frequency in most of the mutants reached 71.8% to 98.9%. The mutants showed 71.2% to 99.8% reduction in citrus canker lesion and greater than 99.7% or 2.45 to 4.95 Log10 unit reduction in Xcc bacterial cell population. Mutants also accumulated more salicylic acid and expressed much higher levels of the *NPR1* gene than the wildtype with or without Xcc inoculation, which suggests potential resistance to other diseases in these mutants. The guide RNAs for targeting *CsDMR6* were selected from highly conserved regions and have 100% nucleotide identity with *DMR6* homologs in important citrus species; these guide RNAs are expected to work in other important citrus species and cultivars.

---

Citrus (*Citrus* L.) is one of the most important fruit crops in the world, grown in more than 114 countries with an acreage of 9.3 million hectares and 146.6 million tons of production (FAO statistics, http://www.fao.org/faostat/en/; Gmitter and Talon, 2008). Sweet oranges (*C*. × *sinensis*), grapefruit (*C*. × *paradisi*), pummelos (pomelos) (*C*. *maxima*), lemons (*C*. × *limon*), limes, mandarins (*C*. *reticulata*), tangerines (*C*. *reticulata*), etc. are the major cultivated types of citrus. Production worldwide faces major challenges from multiple devastating bacterial diseases, including citrus canker, citrus greening (or Huanglongbing, HLB), and citrus variegated chlorosis (CVC). Citrus canker is caused by *Xanthomonas citri* ssp. *citri* (Xcc) (Gottwald and Graham, 2014), and most citrus species and cultivars are susceptible to citrus canker. Improving resistance to citrus canker has been an important citrus breeding objective (Gmitter and Talon, 2008). Recent studies have shown that disabling disease susceptibility (*S*) genes including *DOWNY MILDEW RESISTANCE* 6 (*DMR6*) can be a promising approach to engineering durable and broad-spectrum resistance to bacterial pathogens (de Toledo Thomazella, *et al*., 2021; Hasley, *et al*., 2021; Kieu, *et al*., 2021; Tripathi, *et al*., 2021).

The *DMR6* gene encodes a salicylic acid 5-hydroxylase (Zhang *et al*., 2017) and has been identified as a susceptibility factor to a number of bacterial and oomycete pathogens. DMR6 protein is a repressor of plant immunity (Zeilmaker *et al*., 2015) that negatively regulates the expression of plant defense genes. The expression of *DMR6* is required for plant susceptibility to the downy mildew pathogen *Hyaloperonospora* in *Arabidopsis*, susceptibility to *Pseudomonas syringae* pv. *Tomato* DC3000 (Zhang *et al*., 2017), and susceptibility to the oomycete *Phytophthora capsici* (Van Damme *et al*., 2005). Disruptive mutations of *DMR6* have resulted in resistance to diseases (Zeilmaker *et al*., 2015). CRISPR-Cas9-mediated mutagenesis of a *DMR6* ortholog in tomato resulted in broad-spectrum resistance to *P*. *syringae*, *P*. *capsica*, *Xanthomonas gardneri*, and *X*. *perforans* (del Toledo Thomazella *et al*., 2021). Gene expression studies suggest that *DMR6* may be involved in citrus susceptibility to HLB (Wang *et al*., 2016; Ibanez et al., 2019). Here, we have demonstrated for the first time that CRISPR-Cas9 mediated mutagenesis of a *DMR6* ortholog (*CsDMR6*) in two *Citrus* cultivars, ‘Duncan’ grapefruit (*Citrus* × *paradisi*) and Carrizo citrange (*Citrus* × *sinensis* × *Poncirus trifoliata*), results in strong resistance to citrus canker caused by Xcc.

We confirmed the sequence of a 469-bp region of the coding region of *CsDMR6* in ‘Duncan’ grapefruit and Carrizo citrange and designed two guide RNAs [dmr6-gRNA1 (***CCT***CGGGAATCCGGTACACAAAC), and dmr6-gRNA2 (AGTGGAAAGAGTCTTAGAAG***TGG***)] to target *CsDRM6*. Both guide RNAs have 100% nucleotide identity over their entire length with the *DMR6* homologs in sweet orange (*C*. × *sinensis*), Clementine mandarin (*C*. *clementina*), mandarins (*C*. *reticulata*), lime (*C*. *aurantifolia*), pummelo (*C*. *maxima*), *C*. *ichangensis*, and trifoliate orange (*Poncirus trifoliata*) in the Citrus Genome Database. A plasmid vector (*pD6g1g2-HypaCas9*) (Figure 1A) was constructed to express the two guide RNAs as well as the HypaCas9, GFP and NPTII genes in citrus. The expression of the guide RNAs was driven by the AtU6-26 promoter, and the two guide RNAs were linked using the polycistronic-tRNA-gRNA system. The HypaCas9 and the GFP-NPTII fusion gene were placed after the parsley (*Petroselinum crispum*) Ubiquitin 4-2 promoter (PcUbi4-2) (Nguyen *et al*., 2021) and the enhanced cassava vein mosaic virus promoter, respectively. The vector was introduced into *Agrobacterium tumefaciens* EHA101 strain, and the gene editing cassette was then transferred into citrus through *Agrobacterium* infection and co-cultivation. Co-cultivation of 1900 ‘Duncan’ and 5320 Carrizo epicotyl segments with this *Agrobacterium* strain followed by kanamycin selection and green fluorescence protein (GFP) visualization resulted in identification of nine and 57 GFP-positive shoots, respectively. Based on the GFP expression, the transformation efficiency in ‘Duncan’ and Carrizo was 0.47% and 1.07%, respectively. Micro-grafting of GFP-positive shoots onto Carrizo rootstock seedlings resulted in establishment of four ‘Duncan’ and 16 Carrizo GFP-positive complete plants in soil, of which two ‘Duncan’ and four Carrizo lines were analyzed for induced mutations and resistance to Xcc.

**Figure 1.**
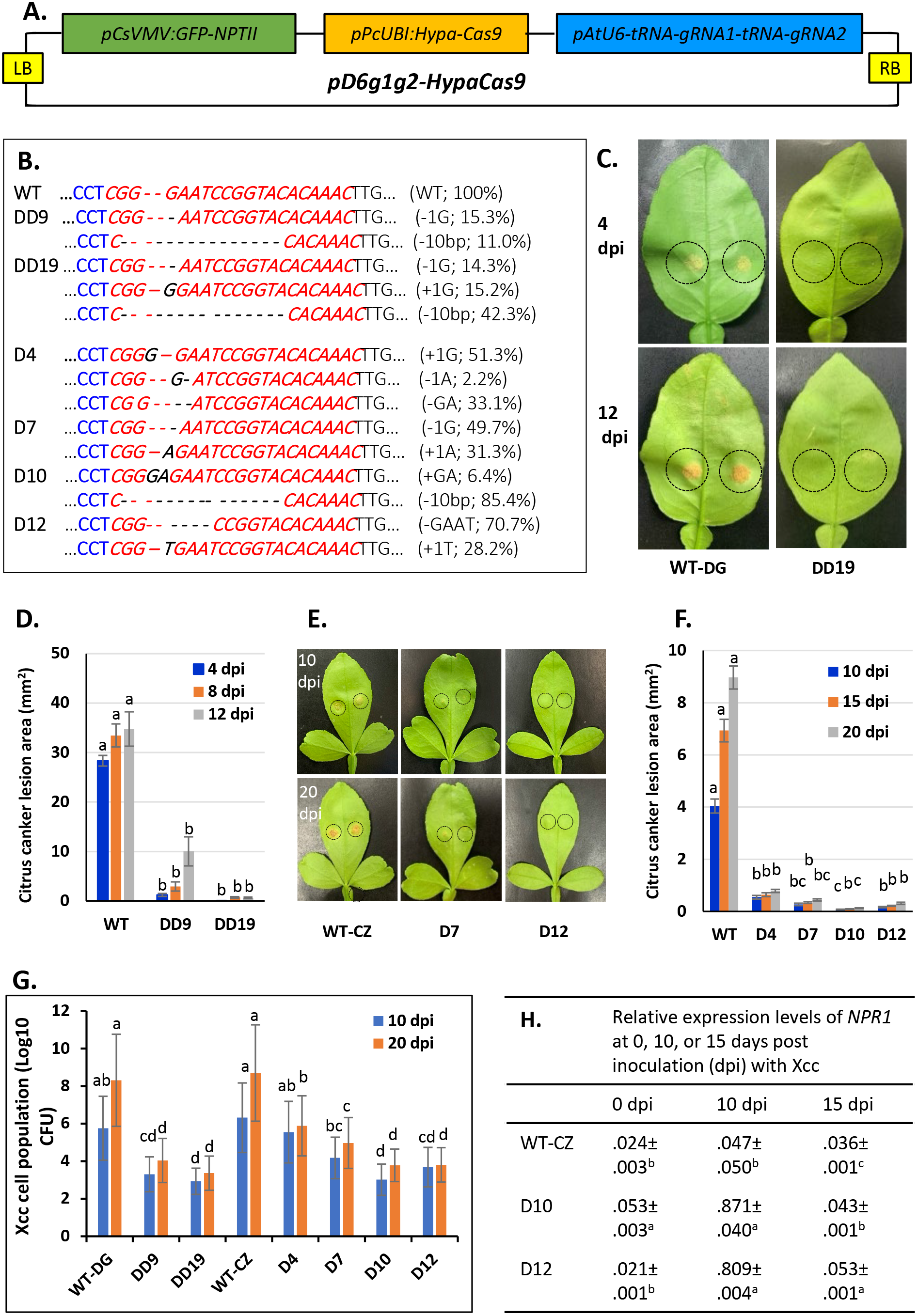
Citrus *dmr6* mutants show resistance to the bacterial disease citrus canker caused by *Xanthomonas citri* ssp. *citri* (Xcc). A. Schematic map of the plasmid vector (*pD6g1g2*-*HypaCas9*) for CRISPR/Cas9-mediated mutagenesis of the *DMR6* ortholog in *Citrus* (*CsDMR6*). B. DNA sequences of wildtype (WT) and six *dmr6* mutants at the first guide RNA-targeted region, mutations, and mutation frequencies (in parenthesis) in ‘Duncan’ grapefruit and Carrizo citrange. C and D. Citrus canker lesions (C) (within the dotted circles) and lesion size (mm^2^) (D) on leaves of wildtype ‘Duncan’ grapefruit (WT-DG) and its *dmr6* mutant (DD19) incited by inoculated Xcc at 4, 8 (in D only), and 12 days post inoculation (dpi). E and F. Citrus canker lesions (E) (within the dotted circles) and lesion size (mm^2^) (F) on leaves of wildtype Carrizo citrange (WT-CZ) and its *dmr6* mutants (D7 and D12) incited by inoculated Xcc at 10, 15 (in F only) and 20 dpi. G. Xcc cell populations (Log10 of colony-forming units per cm^2^) in leaves of wildtype ‘Duncan’ (WT-DG) and Carrizo (WT-CZ) and their *dmr6* mutants (‘Duncan’ DD9 and DD19, and Carrizo D4, D7, D10, and D12) at 10 and 20 dpi. H. Relative expression of the plant defense gene *NPR1* in the leaves of wildtype Carrizo and *dmr6* mutants D10 and D12 prior to and 10 and 15 dpi. The glyceraldehyde 3-phosphate dehydrogenase (*GAPDH*) gene was used as a reference gene to calculate the relative gene expression level of *NPR1* using the 2^−ΔΔ(Ct)^ method.

Deep amplicon sequencing of the targeted *CsDMR6* region revealed multiple types of mutations and different mutation frequencies in ‘Duncan’ (DD9 and DD19) and Carrizo mutant lines (D4, D7, D10, and D12) (Figure 1B). Line DD9 contained two primary types of mutations and exhibited a mutation frequency of 38.5%. Line DD19 contained three primary types of mutations and showed a mutation frequency of 74.2%. The primary mutations in line D4 were one-base insertion and two-base deletion, and this line had a mutation frequency of 89.0%. The primary mutations in line D7 were one-base deletion and one-base insertion, and the mutation frequency was 84.6%. The primary mutation in line D10 was 10-base deletion, and this mutant had a mutation frequency of 91.8%. The primary mutations in line D12 were four-base deletion and seven-base deletion, and D12 showed a mutation frequency of 100%.

Significant differences were observed in the dmr6-gRNA1 and dmr6-gRNA2-targeted regions in term of types and frequencies of mutation. Nine types of mutations were induced in the six mutants in the dmr6-gRNA1-targeted region, including the deletion of one, two, four, or 10-bases, or the insertion of one or two bases of different nucleotides (Figure 1B). These deletions and insertions resulted in frameshifting of the coding region in *CsDMR6*. The mutation frequency in this region in five out of six mutants ranged from 71.8% (DD19) to 98.9% (D12). Fewer types of mutations (three) and much lower frequencies of mutation were induced in the dmr6-gRNA2-targeted region. In five out of the six mutants, the mutation frequency in the dmr6-gRNA2-targeted region was 2.5% to 12.2%, except for line D12, which had a 71.1% mutation frequency.

We inoculated the abaxial side of immature leaves of these mutants and wildtypes with cell suspensions [1 × 10^8^ colony-forming units (CFU)/mL] of Xcc strain 2004-0059, the main strain in Florida, and then measured citrus canker lesions and determined Xcc bacterial cell populations at the inoculated sites. The inoculation experiments were repeated three times, and each inoculation experiment was conducted on multiple leaves. Compared to wildtype ‘Duncan’ grapefruit, mutant DD9 and DD19 showed 71.2% and 99.8% reduction in canker lesion at 4, 8, or 12 days post inoculation (dpi) (Figure 1C, 1D). The Xcc cell counts in DD9 and DD19 leaves at the inoculated sites were reduced by >99.7%, or by 2.45 to 4.95 Log10 units, at 10 or 20 dpi (Figure 1G). Compared to wildtype Carrizo, the citrus canker lesion on leaves of D4, D7, D10, and D12 was reduced by 86.5% to 98.7% at 10, 15, or 20 dpi (Figure 1E, 1F). The Xcc cell population in the leaves of D4, D7, D10, and D12 were reduced by 99.8% or greater, or 2.81 to 4.92 Log10 units at 20 dpi (Figure 1G).

Salicylic acid (SA) accumulation is essential for expression of plant disease resistance (Delaney et al., 1994). *DMR6* gene is known to encode a SA hydroxylase (Zhang *et al*., 2017). We measured SA contents in the leaves of wildtype and mutant ‘Duncan’ grapefruit immediately prior to Xcc inoculation and 24 hours post inoculation (hpi). Prior to Xcc inoculation, mutant line DD9 and DD19 contained 116.2% and 95.0% more SA in the leaves than the wildtype. After Xcc inoculation, these lines showed a rapid increase of SA content and contained 30.1% and 31.0% more SA than the wildtype. The *non-expressor of pathogenesis-related genes 1* (*NPR1*) plays a fundamental role in plant response to pathogen challenge. *NPR1* acts as the master key to the plant defense signaling network. Higher expression levels of *NPR1* are essential for establishing systemic acquired resistance (SAR), as well as induced systemic resistance (ISR). Before *Xcc* inoculation (0 dpi), 15 and 20 dpi, the expression levels of *NPR1* in wildtype Carrizo and mutant lines D10 and D12 were low, ranging from 0.021 to 0.053. However, by 10 dpi, *NPR1* expression increased by 16-39 fold in both mutants whereas it increased by ~2 fold in wildtype (Figure 1H).

In summary, we have demonstrated that disruptive (frameshift) mutagenesis of the *DMR6* homolog (*CsDMR6*) in two citrus cultivars resulted in strong resistance to citrus canker, an important bacterial pathogen to the global citrus industry. We also have shown that the hyper-accurate HypaCas9 mediated high frequencies of mutations in citrus. The guide RNAs designed in this study are expected to target *DMR6* orthologs in multiple commercially important citrus species and cultivars. SA plays a central role in plant resistance to many pathogens. Functional knocking down of *CsDMR6* increased SA accumulation and *NPR1* expression in citrus, thus editing this gene may be an effective approach to improving citrus resistance to other pathogens.

## Acknowledgements

This project was supported by the USDA NIFA Specialty Crop Research Initiative/Citrus Disease Research and Extension (SCRI/CDRE) (2017-70016-26051) grants to F.L., F.G.G., Z.D. and Y.D. The authors thank Dr. Jeff Jones for providing the Xcc strain 2004-00059 and advice on assessing citrus plants for resistance to citrus canker. The authors express appreciation to Mr. Joseph Alexander for providing ‘Duncan’ grapefruit seeds for some of the experiments.

## Author contributions

SP constructed the plasmid vector, generated and identified the mutants, determined the resistance of mutants to citrus canker, and their SA content and gene expression; HH provided entry-level plasmids for CRISPR vector construction; ZD, FGG, YD, and FL conceived the project and secured funding; ZD supervised the project and drafted and finalized the manuscript. All authors reviewed, revised, and approved the manuscript.

